# FRA1 drives melanoma metastasis through an actionable transcriptional network

**DOI:** 10.1101/2025.06.07.658418

**Authors:** Xiaonan Xu, Manon Chadourne, Zulaida Soto-Vargas, Vinesh Jarajapu, Jiqiang Yao, Xiaoqing Yu, Florian A. Karreth

## Abstract

Transcriptional dysregulation has emerged as a critical driver of melanoma progression, yet the molecular mechanisms governing this process and their potential as therapeutic targets remain inadequately characterized. Here, we identify FRA1 as a potent and actionable driver of melanoma metastasis. FRA1 enhanced both the initial seeding and subsequent outgrowth of metastatic lesions. Comprehensive multi-omics integration revealed transcriptional target genes of FRA1, with AXL, CDK6, and FSCN1 exhibiting increased expression in melanoma metastasis and a significant correlation with poor patient outcomes. Silencing AXL, CDK6, or FSCN1 abrogated FRA1-mediated invasion in vitro and reduced metastatic colonization. Furthermore, pharmacological inhibition of CDK6 and FSCN1, and to a lesser extent AXL, suppressed melanoma metastasis and prolonged overall survival. The expression of FRA1 and its target genes correlates with shortened survival across multiple cancer types, highlighting the broader clinical relevance of this pathway. This study unveils an actionable FRA1-mediated transcriptional network that drives cancer progression and metastasis, offering potential avenues for therapeutic interventions.

**SIGNIFICANCE:** FRA1 promotes melanoma metastasis by enhancing the expression of AXL, CDK6, and FSCN1 and this transcriptional network is associated with poor survival across several cancer types. Targeting these FRA1 effectors suppresses metastasis and extends survival, offering a therapeutic strategy for metastatic melanoma and potentially other aggressive cancers.

## INTRODUCTION

Melanoma is an aggressive cancer that affected ∼332,000 people in 2022, with ∼59,000 patients succumbing to the disease worldwide (1). Metastatic melanoma accounts for only 1% of all skin cancer cases but causes the majority of skin cancer-related deaths (2). Despite the recent clinical advances of targeted therapy and immunotherapy, the 5-year survival rate of metastatic melanoma remains poor and additional therapeutic strategies are needed. The development of melanoma is a multi-step process, including melanocyte transformation, nevus formation, malignant transformation, radial growth, vertical growth (invasion), and metastasis. Recurrent mutations, including the activation of oncogenes (BRAF, NRAS) and loss of tumor-suppressors (PTEN, CDKN2A), drive the early stages of melanoma formation (2,3). However, the genetic and molecular events driving melanoma invasion and metastasis remain poorly characterized. No recurrent mutation specifically associated with melanoma metastasis has been identified, and the somatic mutation burden during metastasis remains largely unchanged (2). This indicates that non-genetic events may be critical drivers of melanoma metastasis.

Recent studies highlighted the importance of transcriptional deregulation in many processes involved in cancer metastasis, including epithelial-mesenchymal transition (EMT), invasion, survival in circulation, and colonization of distant organs (4). However, how transcriptional alterations affect melanoma metastasis is largely unknown. Our recent study characterizing the molecular mechanisms underlying PTEN-mediated melanoma suppression identified an AKT/mTORC1/FRA1 pathway that mediates melanoma growth and invasion (5). FRA1, encoded by the FOSL1 gene, belongs to the FOS family and can dimerize with proteins of the JUN family to form the AP-1 transcription factor complex (6). In melanoma, we and others reported that FRA1 is involved in melanoma cell proliferation, anchorage-independent growth, and migration in vitro (5,7). While these findings suggest a role for FRA1 in melanoma progression, its functions specifically in melanoma metastasis have not been elucidated.

In this study, we uncovered that FRA1 robustly promotes melanoma metastasis by enhancing melanoma cell colonization and outgrowth in distant organs. Multi-omics integration defined direct transcriptional targets of FRA1 that are related to pro-metastatic cellular processes and a melanoma metastasis gene signature. Importantly, FRA1 transcriptionally activates *AXL*, *CDK6*, and *FSCN1*, which critically contribute to FRA1-mediated invasion and metastasis. This FRA1-controlled transcriptional network is evident across multiple cancer types and associated with poor outcomes. This study establishes FRA1 as a potent driver of a pro-metastatic, therapeutically actionable transcriptional network in melanoma and other cancers.

## RESULTS

### FRA1 is a driver of melanoma metastasis

We recently identified that PTEN suppresses melanomagenesis in part by opposing AKT/mTORC1-mediated translation of the AP1 transcription factor FRA1 (5). This signaling axis appears to primarily promote melanoma cell invasiveness and xenograft tumor growth, with limited contribution to proliferation (5). FRA1 is associated with an invasive cell state and promotes migration of human melanoma cells in vitro (5). Notably, high FRA1 expression correlates with worse patient survival in the skin cutaneous melanoma dataset from The Cancer Genome Atlas (TCGA) (**Fig. 1A**). Based on these findings, we characterized the functions of FRA1 in melanoma metastasis. We ectopically expressed FRA1 in the 1205Lu human melanoma cell line (Supplementary Fig. S1A), and subsequent subcutaneous injection of these cells in NSG mice showed that FRA1 only moderately increased primary tumor growth (**Fig. 1B**); however, the fast tumor growth precluded analyses of spontaneous metastasis in this model. Similarly, using Doxycycline-inducible CRISPR interference (Dox-CRISPRi) to silence FRA1 in A375 (FRA1 high) and 1205Lu (FRA1 intermediate) human melanoma cells (Supplementary Fig. S1B) resulted in moderately decreased primary tumor growth when these cells were subcutaneously injected into NSG mice (**Fig. 1C, D**). In the 1205Lu Dox-CRISPRi transplant model, Doxycycline administration significantly slowed the growth of primary tumors (**Fig. 1D**), providing sufficient time for spontaneous metastasis to develop. Notably, silencing of FRA1 significantly decreased the formation of spontaneous lung metastasis (**Fig. 1E**). To directly assess the effect of FRA1 on metastasis and exclude differences in primary tumor growth as the cause for differential metastasis, we inoculated luciferase-labeled melanoma cells intravenously as an experimental metastasis model. Bioluminescence in vivo imaging showed that FRA1 silencing reduced the metastatic burden of 1205Lu cells in the lung (**Fig. 1F**), while FRA1 overexpression enhanced lung metastasis of 1205Lu melanoma cells (**Fig. 1G**). Tail vein injection of murine M10M6 melanoma cells overexpressing FRA1 (Supplementary Fig. S1A) similarly resulted in enhanced lung metastasis (**Fig. 1H**), indicating that the pro-metastatic effects of FRA1 are conserved across species. H&E staining corroborated the differential lung metastasis burden upon FRA1 silencing and overexpression (Supplementary Fig. S1C-E). Moreover, analysis of a skin cutaneous melanoma TCGA dataset revealed a significant correlation of FRA1 expression with a melanoma metastasis signature (**Fig. 1I**). These findings identified FRA1 as a metastatic driver in melanoma.

**Figure 1.**
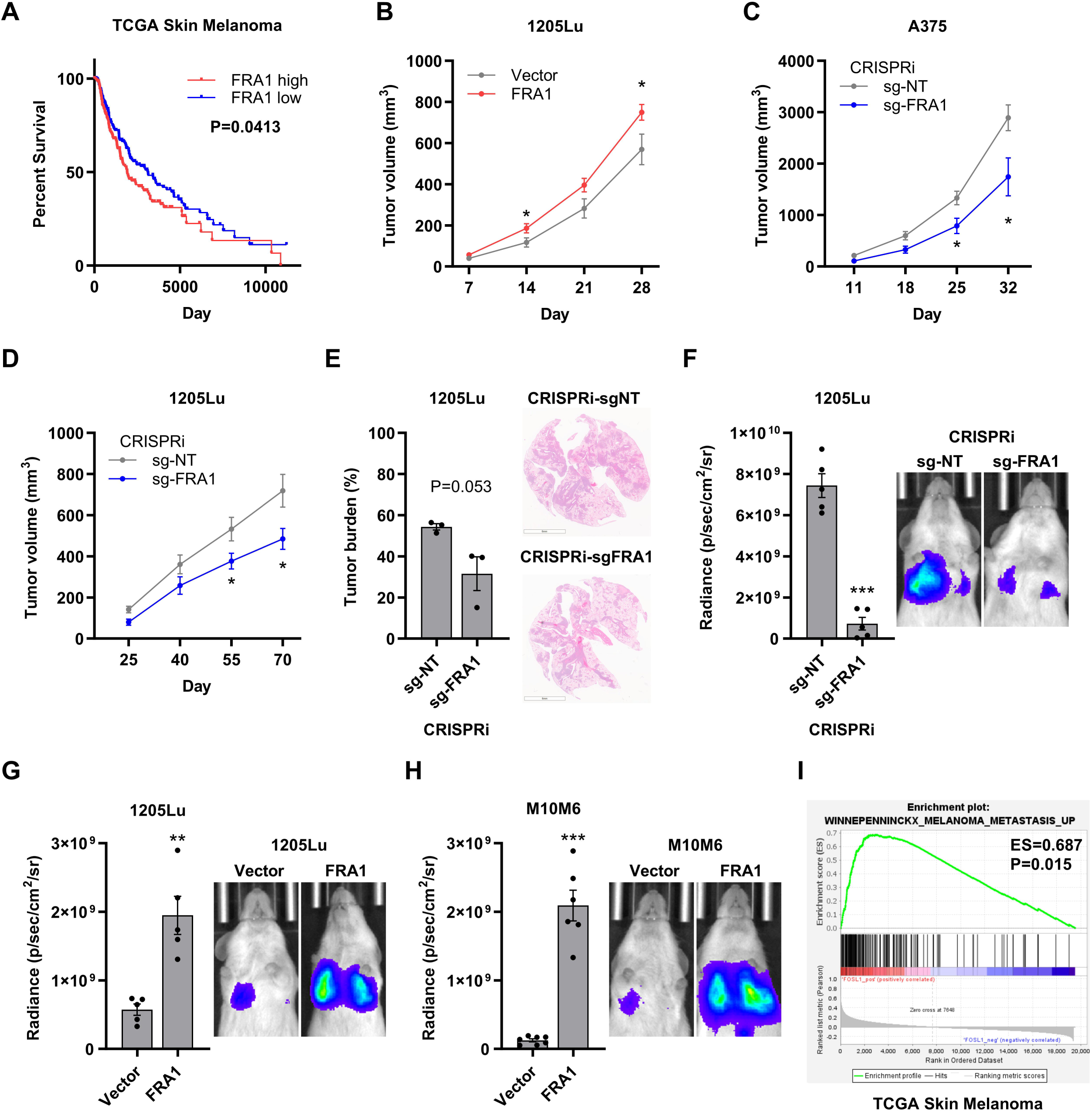
FRA1 drives melanoma metastasis. **A,** Overall survival curve of melanoma patients with high or low FRA1 expression from TCGA. **B,** 1205Lu melanoma cells overexpressing FRA1 were subcutaneously injected into NSG mice (n = 10). Tumor volumes were measured every 7 days. **C-E,** A375 and 1205Lu melanoma cells with inducible CRISPR interference (CRISPRi) targeting FRA1 were subcutaneously injected into NSG mice (n = 6). Mice were fed chow containing 200 mg/kg Dox to induce CRISPRi. Tumor volumes of A375 cell were measured every 7 days (**C**), and tumor volumes of 1205Lu cells were measured every 15 days (**D**). At the endpoint, mice were dissected for detecting spontaneous metastasis. The lung metastasis burden of 1205Lu cells by H&E staining was shown (**E**). **F-H,** Luciferase tagged 1205Lu cells with inducible CRISPRi targeting FRA1 were intravenously injected into NSG mice (n = 5) and the metastases were measured after 22 days by bioluminescent In Vivo Imaging System (IVIS) (**F**). Luciferase tagged 1205Lu and M10M6 cells with FRA1 overexpression were intravenously injected into NSG mice (n = 5), and mice were fed chow containing 200 mg/kg Dox to induce CRISPRi. The metastases were measured after 18 days by IVIS (**G** and **H**). Representative bioluminescence images and quantification of the luminescence signals are shown. **I,** GSEA analysis of TCGA-SKCM samples associates FRA1 expression with a melanoma metastasis gene signature. Data are presented as mean ± SEM and analyzed with Student’s unpaired t test, * P<0.05, ** P<0.01, *** P<0.001.

### FRA1 promotes metastatic colonization and outgrowth

We next determined which stage of the metastatic cascade is influenced by FRA1. To evaluate cell seeding and colonization ability, we performed tail vein injections of 1205Lu melanoma cells and analyzed the formation of micro-metastasis after 7 days. At this time point, micro-metastasis in the lung were comprised of 5-30 cells and their quantification revealed that FRA1 overexpression increased the number of nodules but had no significant effect on nodule size (**Fig. 2A**). We next performed tail vein injections of 1205Lu and A375 cells in which FRA1 transcription was suppressed by Dox-CRISPRi. Silencing of FRA1 significantly decreased the number of micro-metastatic nodules in the lung, with a tendency of also decreasing colony size (**Fig. 2B, C**). Next, we tested the effects of FRA1 on metastatic outgrowth. To this end, we performed tail vein injection of A375 and 1205Lu cells, induced FRA1 silencing through CRISPRi by Dox administration 10 days post-injection, and analyzed metastasis after two weeks on day 24 (**Fig. 2D**). In vivo bioluminescence imaging showed that silencing FRA1 decreased the luciferase signal of 1205Lu cells in the lung (**Fig. 2E**), and H&E staining corroborated the decreased lung metastasis burden upon FRA1 silencing (**Fig. 2F**). Conversely, FRA1 silencing had no significant effect on the outgrowth of A375 cells in the lung (**Fig. 2G**). Because A375 cells have a propensity to metastasize to the liver, we also examined the effect of FRA1 silencing on liver metastasis. Metastatic seeding of A375 cells was difficult to assess as only 2 or fewer nodules were present per liver section at the 7-day timepoint (Supplementary Fig. S1F). However, when FRA1 was silenced starting at day 10 post-injection, the number of liver metastasis and the liver metastatic burden were significantly reduced (**Fig. 2H**). These findings indicate that FRA1 promotes both colonization and outgrowth of metastatic melanoma nodules.

**Figure 2.**
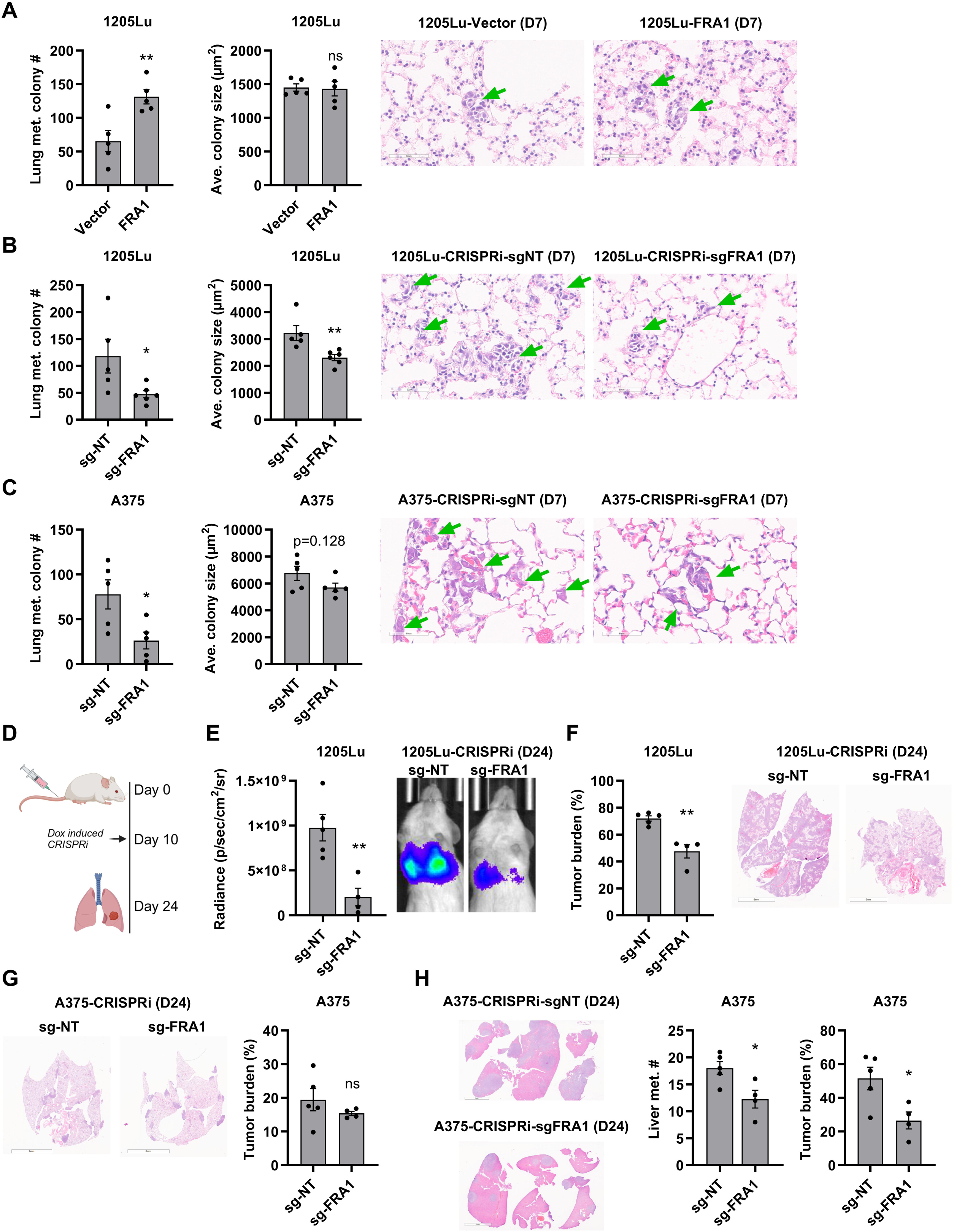
FRA1 enhances metastatic colonization and outgrowth. **A-C,** 1205Lu melanoma cells overexpressing FRA1 were intravenously injected into NSG mice (n = 5) (**A**). 1205Lu and A375 cells with inducible CRISPRi targeting FRA1 were intravenously injected into NSG mice (n = 5), and mice were fed chow containing 200 mg/kg Dox to induce CRISPRi (**B** and **C**). 7 days after inoculation, metastasis was detected by H&E staining. Representative images of micro-metastasis in lungs and quantification of the number and average size of micro-metastasis were shown. **D-H,** Luciferase tagged 1205Lu and A375 cells with inducible CRISPRi targeting FRA1 were intravenously injected into NSG mice (n = 5), and 10 days after inoculation mice were switched to chow containing 200 mg/kg Dox to induce CRISPRi, and then metastasis were analyzed 2 weeks after Dox diet feeding, which is 24 days after cell inoculation (**D**). Representative images and quantification of the luminescence signals of 1205Lu cells are shown in **E**. Representative images and quantification of lung metastasis burden of 1205Lu and A375 cells by H&E staining were shown in **F** and **G**. Representative images and quantification of liver metastasis number and burden of A375 cells were shown in **H**. Data are presented as mean ± SEM and analyzed with Student’s unpaired t test, * P<0.05, ** P<0.01, *** P<0.001, ns not significant.

### FRA1 regulates melanoma metastasis-associated genes

To characterize the mechanism whereby FRA1 promotes melanoma metastasis, we evaluated the transcriptome of 1205Lu and A375 melanoma cells upon FRA1 silencing by RNA sequencing. 590 and 290 differentially expressed protein coding genes were identified in A375 and 1205Lu cells, respectively (**Fig. 3A, B**). The expression changes in 1205Lu and A375 cells upon FRA1 silencing were in concordance and positively correlated (**Fig. 3C**). 52 genes including *FSCN1*, *CLTA*, *RELL1*, and *KIAA0040* were upregulated, while 21 genes including *ATOH8* and *GDF15* were downregulated by FRA1 in both cell lines (**Fig. 3G**, Supplementary Fig. S2A). Gene Set Enrichment Analysis (GSEA) of Hallmark Gene Sets (8) revealed the enrichment of similar gene signatures in A375 and 1205Lu cells upon FRA1 silencing. Specifically, FRA1 expression enhanced signatures associated with Myc targets, E2F targets, G2M checkpoint, mitotic spindle, and DNA repair, and repressed signatures associated with the P53 pathway, apoptosis, interferon response, IL-6 signaling, and TGFβ signaling (**Fig. 3D, E**). Furthermore, GSEA of KEGG pathways demonstrated that FRA1 is associated with cell cycle, DNA replication, and DNA repair signatures, while apoptosis, P53 pathway, focal adhesion, gap junction, JAK-STAT signaling, and TGFβ signaling signatures were depleted (Supplementary Fig. S2B, C). Importantly, GSEA highlighted that FRA1 promotes the enrichment of a melanoma metastasis expression signature (**Fig. 3F**). Taken together, FRA1 regulates genes related to the cell cycle, DNA repair, apoptosis, and induces a melanoma metastasis signature.

**Figure 3.**
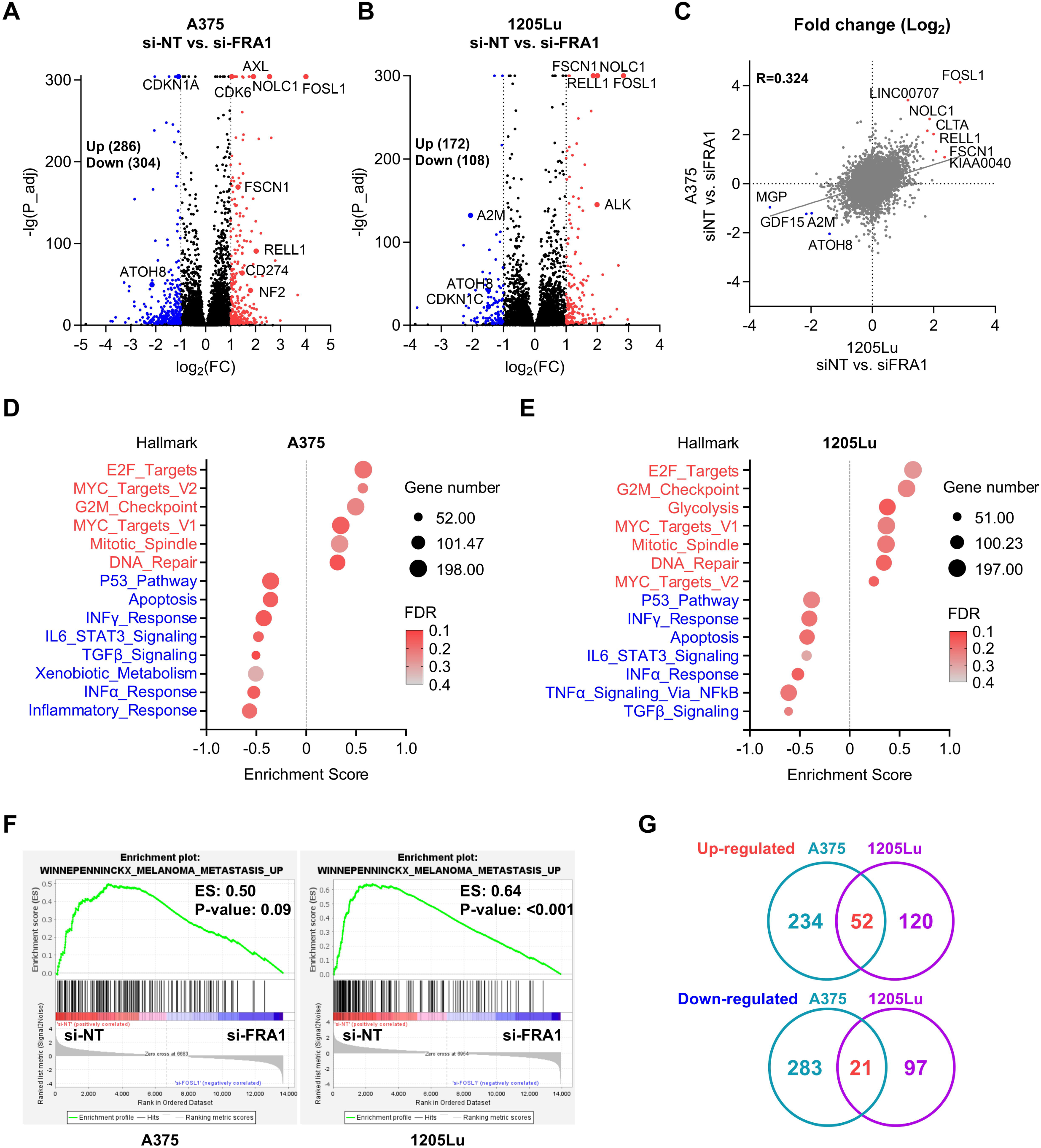
Transcriptome of FRA1 in melanoma. **A-C,** A375 and 1205Lu cells with FRA1 silenced by siRNAs were subjected to RNA sequencing. Volcano plots show the differential expressing genes by FRA1 in A375 cells (**A**) and 1205Lu cells (**B**). Scatter plot shows correlation of gene regulation by FRA1 in A375 and 1205Lu cells (**C**). **D, E,** GSEA analysis of Hallmark gene signature show the enrichment of genes regulated by FRA1 in A375 cells (**D**) and 1205Lu cells (**E**). **F,** GSEA analysis show transcriptome of FRA1 is associated with a melanoma metastasis gene signature. **G,** Overlapped differential expressed genes by FRA1 in A375 and 1205Lu cells.

### Genome-wide identification of FRA1 binding in melanoma

To characterize the genome binding site occupancy of FRA1 in melanoma, we performed Cut&Run followed by sequencing in A375 and 1205Lu cells. 36,576 and 11,671 FRA1 peaks were identified in A375 and 1205Lu cells, respectively, and 7,710 of these FRA1 peaks were shared between both cell lines (**Fig. 4A**). De novo motif analysis of FRA1 genome binding sites identified significant enrichment of the canonical FOS/JUN motif as well as motifs for the TEAD, ETS, and RUNX transcription factor families, indicating that FRA1 may regulate transcription in melanoma by interacting with these transcription factors (**Fig. 4B**). Annotation of the FRA1 occupancy profile revealed that approximately one third of peaks in A375 and 1205Lu cells localized to promoter regions and another third of peaks was intronic (Supplementary Fig. S2D, E). Among the 7,710 FRA1 peaks shared between A375 and 1205Lu cells, 3,267 peaks (42%) localized to promoter regions of 2,882 genes, 2,824 peaks (37%) were within gene bodies (5’UTRs, exons, introns, 3’UTRs), and 1,619 peaks (21%) were in intergenic regions (**Fig. 4C**). The enrichment score of the 3,267 shared promoter peaks strongly correlated between A375 and 1205Lu cells (**Fig. 4D**), indicating a common target gene profile of 2,882 genes, of which 2,143 are protein coding genes. Gene ontology (GO) analysis of biological processes revealed that FRA1 target genes are associated with PDGF signaling, MAPK signaling, apoptosis, DNA damage response, and cell migration and adhesion (**Fig. 4E**), while GO analysis of molecular functions identified an association with kinesin binding, cyclin binding, extracellular matrix (ECM) binding, and cell adhesion molecular binding (**Fig. 4F**). Thus, multi-omics analysis demonstrated that FRA1 regulates genes related to the cell cycle, cell survival, and cell migration and adhesion in melanoma cells.

**Figure 4.**
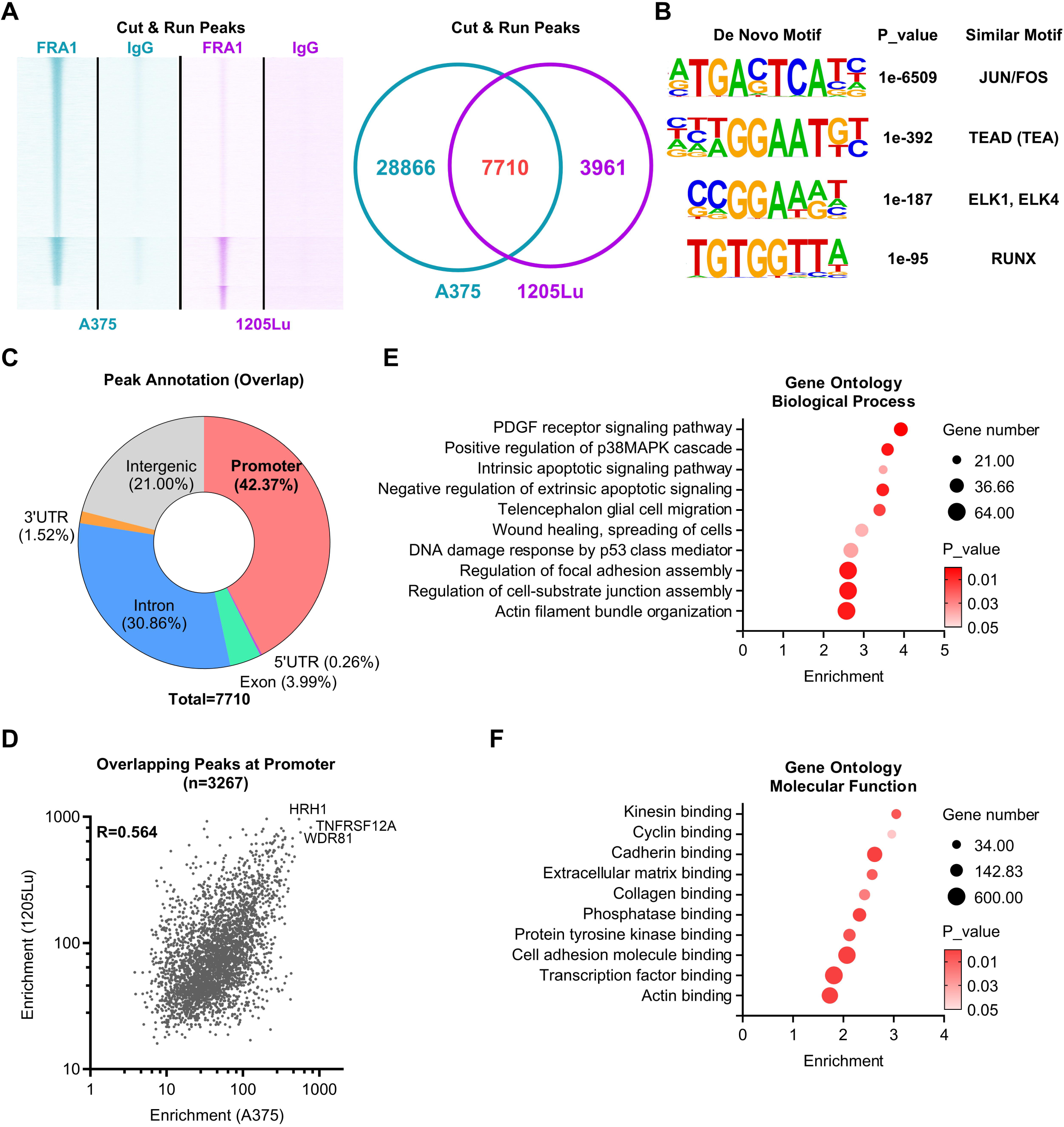
Genome-wide binding profile of FRA1 in melanoma. **A,** A375 and 1205Lu cells were subjected to Cut&Run sequencing by FRA1 and IgG as negative control. Heatmap show the read density of FRA1 on genome in A375 cells and 1205Lu cells. Venn diagram show the overlapped peaks. **B,** De Novo Motif analysis by HOMER show enriched motifs of FRA1 in melanoma cells. **C,** Genomic annotation of overlapped FRA1 bound peaks in A375 and 1205Lu cells. **D,** Scatter plot show the enrichment of overlapping FRA1 peaks in A375 cells and 1205Lu cells. **E, F,** Gene Ontology (GO) analysis of overlapping FRA1 peaks show the enriched gene signature in GO Biological Process (**E**) and GO Molecular Function (**F**).

### FRA1 transcriptionally activates AXL, CDK6, and FSCN1

To identify FRA1 transcriptional targets in melanoma, we integrated the 807 differentially expressing genes identified by RNA-seq and the 2,882 genes demonstrating FRA1 promoter occupancy by Cut&Run-seq. 74 and 48 genes whose promoters showed FRA1 occupancy were upregulated and downregulated, respectively, indicating direct transcriptional regulation by FRA1 (Supplementary Fig. S3A). Moreover, 25 out of 75 genes transcriptionally upregulated by FRA1 exhibit increased abundance in metastasis compared to primary melanoma in the skin cutaneous melanoma TCGA dataset, indicating that they could mediate the pro-metastatic effects of FRA1 (Supplementary Fig. S3B). This included three direct FRA1 targets, *AXL*, *CDK6*, and *FSCN1* (**Fig. 5A**, Supplementary Fig. S3C), that encode proteins (AXL, CDK6, and Fascin) for which pharmacological inhibitors are available. Analysis of the melanoma TCGA dataset revealed that concurrent alterations of *AXL*, *CDK6*, and *FSCN1* correlate with poor outcomes of melanoma patients (**Fig. 5B**). Through modulation of FRA1 expression, we validated at the mRNA and protein levels that FRA1 is a positive regulator of AXL, CDK6, and Fascin (**Fig. 5C**, Supplementary Fig. S3D). Similar to the increase of FRA1 in melanoma cells compared to wildtype or BRAF-mutant melanocytes, AXL, CDK6, and Fascin levels are increased in melanoma cells, especially in cells exhibiting high levels of FRA1 (**Fig. 5D**). Multiplex IHC analysis of a melanoma tumor microarray (TMA) demonstrated that FRA1 is highly expressed in melanoma tissues (**Fig. 5E**). Moreover, 15 out of 19 (79%) lymph node metastatic samples are FRA1 positive, while 22 out of 35 (63%) primary cutaneous melanoma samples are FRA1 positive (**Fig. 5F**). Similarly, more Fascin positive samples were found in metastasis (10 of 19) compared to primary tumors (6 out of 35), while AXL and CDK6 are only positive in 11.1% and 18.5% of all samples, respectively (Supplementary Fig. S3E). Cells in the TMA expressing FRA1 and AXL, CDK6, or Fascin demonstrated a positive correlation of the fluorescent intensity of FRA1 and its targets (**Fig. 5G**), indicating that increased FRA1 expression elevates the levels of AXL, CDK6, and Fascin. Additionally, analysis of DepMap datasets demonstrated that *FRA1* levels significantly correlate with *AXL*, *CDK6*, and *FSCN1* expression in 1,673 cell lines of all cancer types (**Fig. 5H**) and 110 melanoma cell lines (Supplementary S4A) and that FRA1 protein expression also correlates with AXL, CDK6, and Fascin levels in 20 melanoma cell lines (Supplementary S4B). Analysis of the Pan-Cancer dataset integrating International Cancer Genome Consortium (ICGC) and The Cancer Genome Atlas (TCGA) (9) further revealed a significant correlation between *FRA1* levels and *AXL*, *CDK6*, and *FSCN1* expression in 1,210 tumor samples of all cancer types (**Fig. 5I**). These findings indicate that FRA1 directly regulates the expression of *AXL*, *CDK6*, and *FSCN1*.

**Figure 5.**
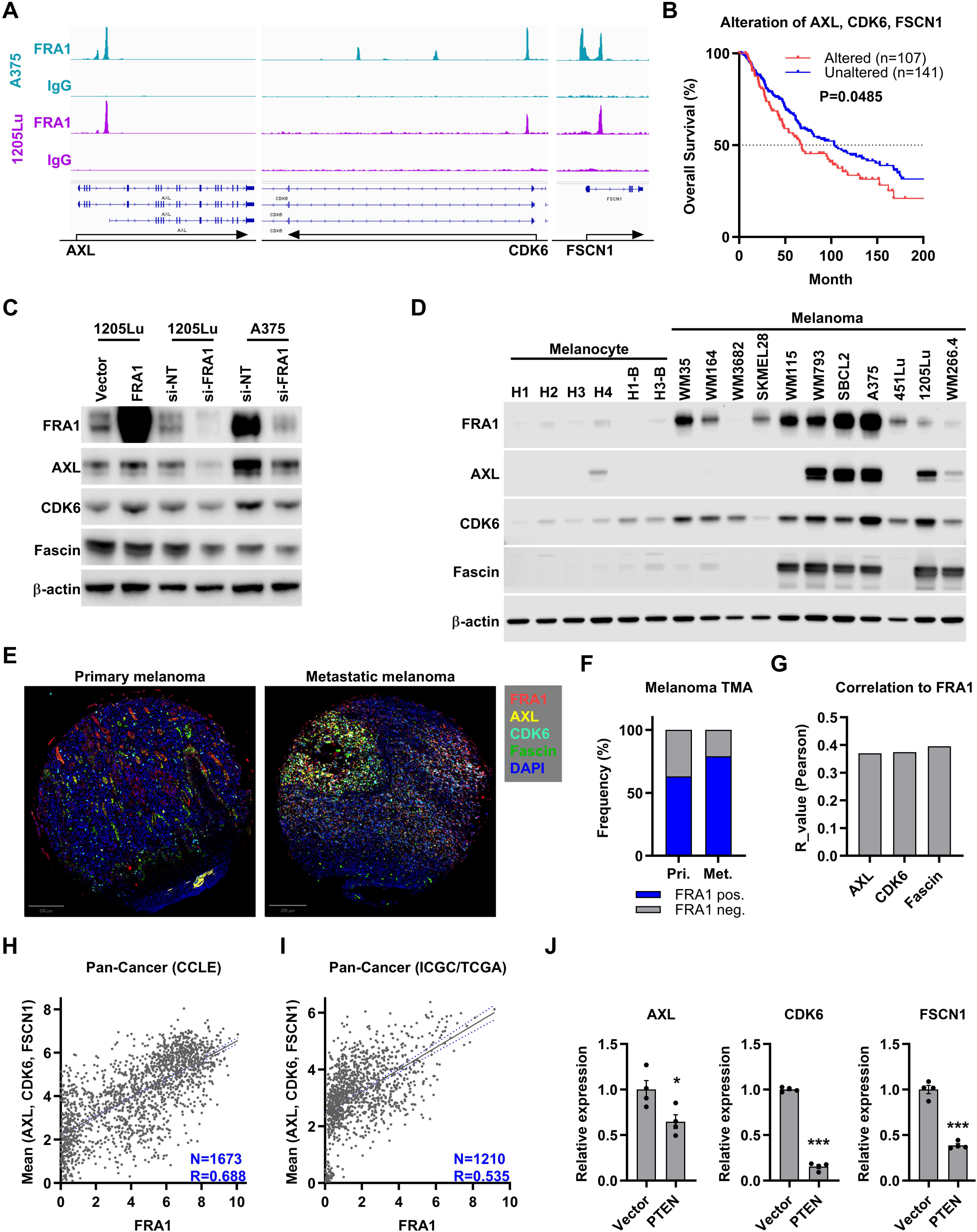
FRA1 transcriptionally activates *AXL*, *CDK6*, and *FSCN1*. **A,** IGV genome tracks highlight FRA1 occupancy at the *AXL*, *CDK6*, and *FSCN1* genomic loci. **B,** Overall survival curves of melanoma patients with or without alteration of *AXL*, *CDK6*, or *FSCN1* (including mutation, copy number alteration, and mRNA changes) in TCGA-SKCM dataset. **C,** Western blot analysis of AXL, CDK6, and Fascin upon FRA1 overexpression or silencing in melanoma cells. **D,** Western blot analysis of FRA1, AXL, CDK6, and Fascin in human melanocytes (H1, H2, H3, H4), human melanocytes expressing oncogenic BRAF^V600E^ (H1-B, H3-B), and melanoma cell lines. **E,** Representative image of multiplex IHC showing the expression of FRA1, AXL, CDK6, and Fascin in a melanoma microarray containing primary melanoma and metastatic melanoma. **F,** Bar plot showing frequency of FRA1 positive samples in primary skin melanomas and lymph node metastatic melanomas. **G,** Pearson correlation analysis of fluorescent intensity of FRA1 and AXL, CDK6, or Fascin. **H** and **I,** Scatter plot showing the correlation between FRA1 expression and the mean expression of *AXL*, *CDK6*, *FSCN1* in 1,673 cancer cell lines (**H,** Data extracted from DepMap) and 1,210 tumor tissues (**I,** Data extracted from cBioPortal). Expression levels are presented as log_2_(TPM+1). **J,** qRT-PCR showing the relative expression of *AXL*, *CDK6*, and *FSCN1* in 1205Lu cells upon expression of PTEN. Data are presented as mean ± SEM and analyzed with Student’s unpaired t test, * P<0.05, ** P<0.01, *** P<0.001.

We previously discovered that FRA1 is translationally regulated by PTEN/AKT/mTORC1 signaling (5), and we therefore determined the impact of PTEN/AKT/mTORC1 signaling on *AXL*, *CDK6*, and *FSCN1* expression. Suppressing the AKT-mTORC1 axis by PTEN overexpression in 1205Lu cells decreased the expression of *AXL*, *CDK6*, and *FSCN1* (**Fig. 5J**). Similarly, the AKT inhibitor MK2206 or the mTOR inhibitor Rapamycin decreased *AXL*, *CDK6*, and *FSCN1* levels in 1205Lu cells (Supplementary Fig. S4C). These data demonstrate that *AXL*, *CDK6*, and *FSCN1* are transcriptionally activated by FRA1 and regulated by PTEN/AKT/mTORC1 signaling.

### AXL, CDK6, and FSCN1 mediate pro-metastatic effects of FRA1 and are potential therapeutic targets in melanoma

We next examined whether FRA1 promotes melanoma metastasis via transcriptionally activating AXL, CDK6, and Fascin expression. To this end, we silenced *AXL*, *CDK6*, or *FSCN1* in FRA1 overexpressing 1205Lu cells. Silencing *CDK6* and *FSCN1*, but not *AXL*, significantly reduced melanoma cell proliferation (**Fig. 6A**). Moreover, silencing *AXL*, *CDK6*, or *FSCN1* significantly reversed the stimulation of melanoma cell invasion by FRA1 (**Fig. 6B**). To evaluate the effects of AXL, CDK6, and Fascin on cell colonization during distant metastasis, we intravenously inoculated 1205Lu cells transfected with siRNAs in NSG mice and analyzed the formation of micro-metastasis after 7 days. Silencing *AXL*, *CDK6*, and *FSCN1* completely reverted the FRA1-induced increase in the number of micro-metastatic nodules while the average colony sizes remained unchanged (**Fig. 6C**). To determine whether AXL, CDK6, and Fascin affect metastatic outgrowth, we intravenously inoculated luciferase-tagged 1205Lu cells and monitored metastasis by in vivo bioluminescence imaging. 14 days after injection when we first detected bioluminescent signal from lung metastasis, we started the administration of the AXL inhibitor R428, the CDK4/6 inhibitor G1T38, or the Fascin inhibitor NP-G2-044 every other day for four treatments (**Fig. 6D**). The CDK4/6 and Fascin inhibitors, and to a lesser extent the AXL inhibitor, decreased lung metastasis burden compared to vehicle control (**Fig. 6E**; Supplementary Fig. S4D), indicating that FRA1 promotes metastatic outgrowth via its targets *AXL*, *CDK6*, and *FSCN1*. To determine whether the combination of CDK4/6 and Fascin inhibitors exerts enhanced therapeutic efficacy, we treated NSG mice inoculated with 1205Lu cells daily with the CDK4/6 inhibitor Abemaciclib and the Fascin inhibitor NP-G2-044 (**Fig. 6F**). Notably, the median overall survival of the cohort receiving the combination treatment was significantly extended compared to the vehicle control cohort (27 vs. 18 days) (**Fig. 6G**). CT scans on days 10 and 18 showed decreased lung metastatic burden in the combination treatment cohort compared to the vehicle control group (**Fig. 6H**). Importantly, no significant toxicity or adverse drug reactions were observed during the treatment. These findings demonstrate that FRA1 enhances melanoma metastasis at least in part by transcriptionally activating *AXL*, *CDK6*, and *FSCN1*, and targeting CDK4/6 and Fascin could be explored as a therapeutic strategy for FRA1-high metastatic melanoma.

**Figure 6.**
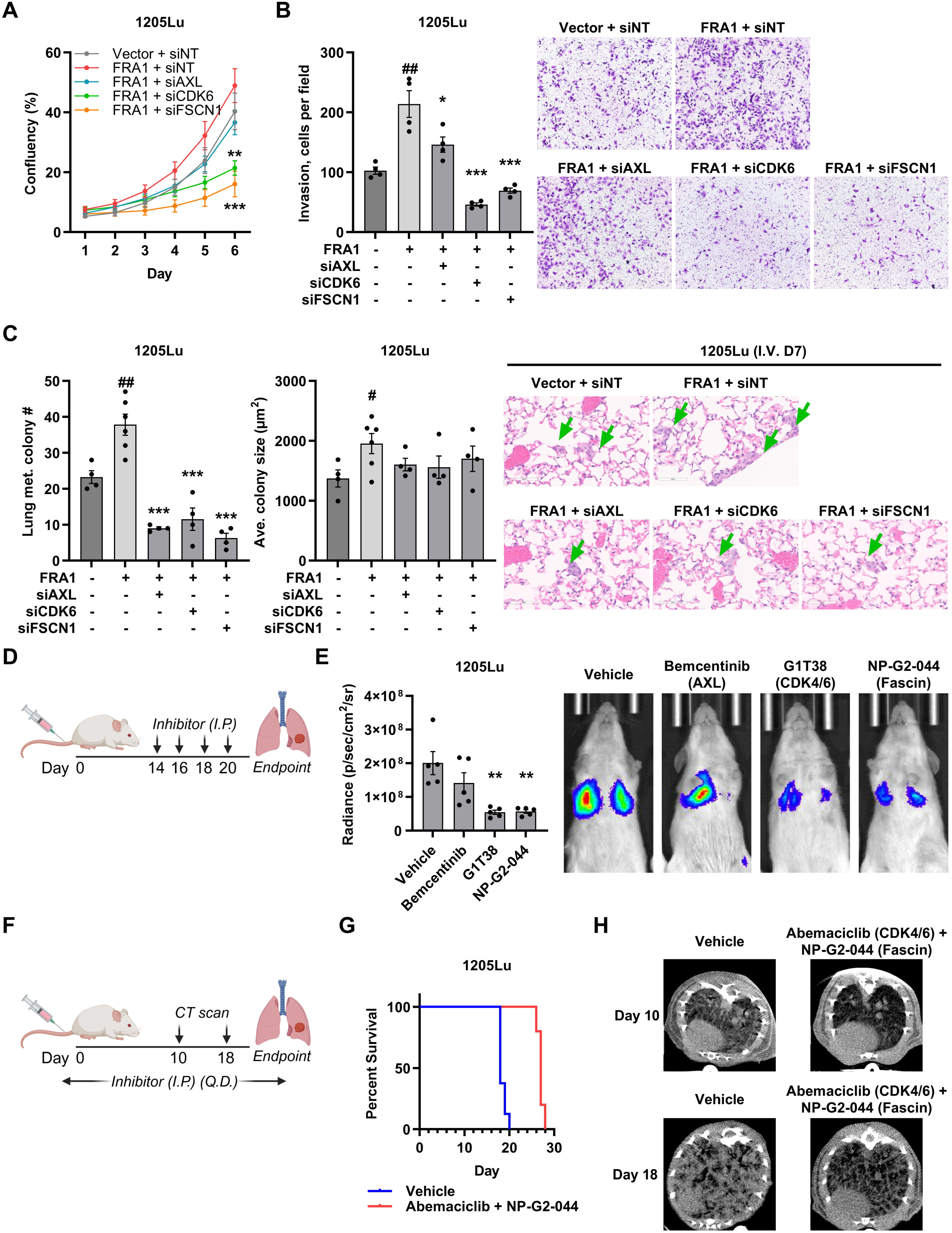
AXL, CDK6, and FSCN1 mediate FRA1 pro-metastatic functions. **A,** Proliferation as measured by relative confluence of 1205Lu cells expressing FRA1 with AXL, CDK6, or FSCN1 silenced. **B,** Representative images and quantification of cell numbers of transwell invasion assays of 1205Lu cells expressing FRA1 with AXL, CDK6, or FSCN1 silenced. **C,** FRA1 overexpressing 1205Lu cells with AXL, CDK6, or FSCN1 silenced were intravenously injected into NSG mice (n = 5). 7 days after inoculation, metastasis was detected by H&E staining. Representative images of micro-metastasis in lungs and quantification of the number and average size of micro-metastasis were shown. **D, E,** Luciferase tagged 1205Lu cells were intravenously injected into NSG mice. AXL inhibitor Bemcentinib, CDK4/6 inhibitor G1T38, Fascin inhibitor NP-G2-044, and vehicle control were intraperitoneally (I.P.) administrated to mice (n = 5) at day 14, 16, 18, and 20 after cell inoculation (**D**). Metastasis was measured at day 20 by IVIS, and representative images and quantification of luminescence signals were shown (**E**). **F-H,** 1205Lu cells were intravenously injected into NSG mice. Combination of CDK4/6 inhibitor Abemaciclib and Fascin inhibitor NP-G2-044, or vehicle control were intraperitoneally (I.P.) administrated to mice (n=5) once per day (Q.D.) until endpoint (**F**). Overall survival curves of mice with combo treatment or vehicle control were shown (**G**). Metastasis were detected by CT scan at day 10 and 18, representative images were shown (**H**). Data are presented as mean ± SEM and analyzed with Student’s unpaired t test, * P<0.05, ** P<0.01, *** P<0.001.

### Evidence for oncogenic FRA1 signaling across multiple cancer types

FRA1 upregulation has been reported in multiple cancer types and is associated with cancer progression (6). To investigate the effect of FRA1 upregulation on patient outcomes across cancer types, we analyzed TCGA’s Pan Cancer Atlas. High *FOSL1* expression is associated with poor overall survival in lung adenocarcinoma, pancreatic adenocarcinoma, clear cell renal cell carcinoma, sarcoma, low grade glioma, glioblastoma, head and neck squamous carcinoma (HNSCC), and uveal melanoma (**Fig. 7A**). Moreover, the concurrent alteration of the FRA1 targets, *AXL*, *CDK6*, and *FSCN1* correlated with worse prognosis in lung adenocarcinoma, low grade glioma, and clear cell renal cell carcinoma (Supplementary Fig. S5A). Analysis of TCGA’s Pan Cancer Atlas as well as the Cancer Cell Line Encyclopedia (CCLE) dataset revealed robust correlations between *FOSL1* expression and *AXL*, *CDK6*, or *FSCN1* levels in tissues of multiple cancers, especially cancers of the lungs, liver, bladder, pancreas, colon, breast, and head and neck (**Fig. 7B,C**). These findings suggest that the FRA1-regulated transcriptional network contributes to the progression in multiple cancer types.

**Figure 7.**
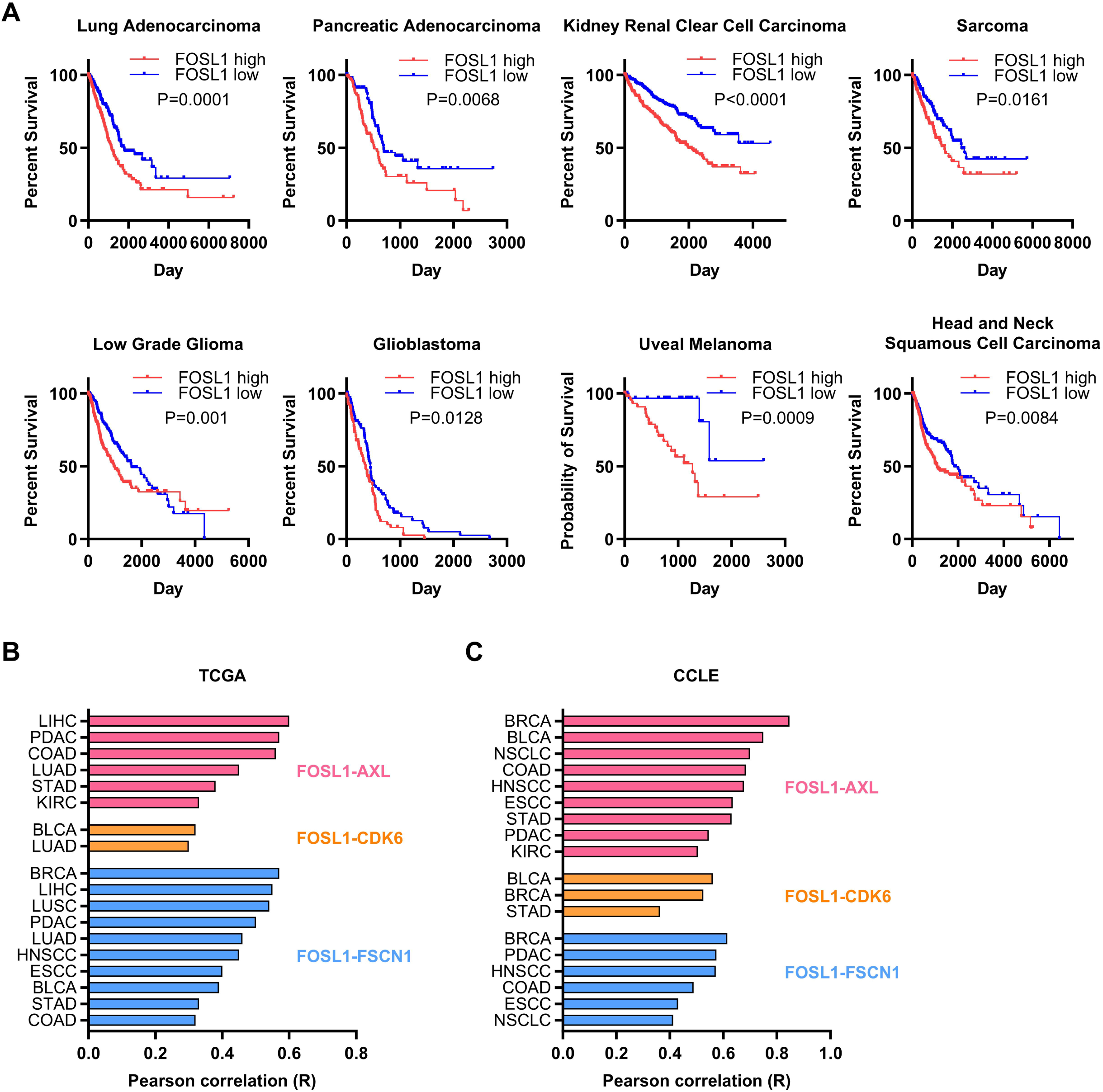
Evidence of oncogenic FRA1 signaling in multiple cancer types. **A,** Overall survival curve of patients with high or low FRA1 expression in multiple cancer types from TCGA Pan Cancer Atlas dataset. **B,** Correlation between FRA1 expression and expression of AXL, CDK6, FSCN1 in multiple cancer types from TCGA Pan Cancer Atlas dataset. **C,** Correlation between FRA1 expression and expression of AXL, CDK6, FSCN1 in multiple cancer types from Cancer Cell Line Encyclopedia (CCLE) (Data extracted from DepMap). (LIHC: Liver hepatocellular carcinoma. PDAC: Prostate adenocarcinoma. CODA: Colon adenocarcinoma. LUAD: Lung adenocarcinoma. LUSC: Lung squamous cell carcinoma. NSCLC: Non-small cell lung cancer. STAD: Stomach adenocarcinoma. KIRC: Kidney renal clear cell carcinoma. BLCA: Bladder urothelial carcinoma. BRCA: Breast invasive carcinoma. HNSCC: Head and neck squamous cell carcinoma. ESCC: Esophageal carcinoma.)

## DISCUSSION

This study identifies FRA1 as a critical driver of melanoma metastasis, revealing its role in enhancing both colonization and outgrowth of metastatic lesions. While genetic mutations such as BRAF activation and PTEN loss drive early melanoma formation, the mechanisms underlying metastasis remain less characterized. Our findings demonstrate that transcriptional dysregulation through FRA1 provides a crucial non-genetic mechanism promoting metastatic progression of melanoma, expanding our current knowledge of transcriptional control of metastasis (10).

FRA1 specifically enhances the ability of melanoma cells to colonize distant organs and promotes the outgrowth of micro-metastases, processes considered rate-limiting in the metastatic cascade. We demonstrated that FRA1 silencing significantly reduced both the initial seeding and subsequent growth of metastatic nodules. Notably, FRA1 promotes metastasis across different organ microenvironments, as evidenced by its effects in both lung-tropic 1205Lu cells and A375 cells that metastasize to multiple organs, suggesting a broadly applicable pro-metastatic mechanism independent of organ-specific factors.

Prior studies have uncovered roles for FRA1 in melanomagenesis. FRA1 has been implicated in melanocyte transformation, where it cooperates with BRAF^V600E^ to promote cellular transformation and proliferation (11). FRA1 also contributes to therapy resistance in melanoma, with elevated FRA1 levels observed in BRAF inhibitor-resistant cells (12,13). In treatment naïve melanoma cells, FRA1 regulates therapy-induced secretomes, influencing the tumor microenvironment and promoting adaptive resistance to targeted therapies (14). While these studies established the role of FRA1 in melanoma initiation and therapy resistance, our work specifically characterizes its critical functions in metastatic progression. Collectively, these studies highlight the prominent and context-specific functions of FRA1 in melanoma biology.

Through multi-omics integration, we identified a FRA1-regulated transcriptional network controlling cell cycle progression, DNA repair, apoptosis resistance, and cell migration/adhesion. GSEA revealed that FRA1 enhances signatures associated with MYC targets, E2F targets, G2M checkpoint, and DNA repair, while repressing signatures related to p53 pathway, apoptosis, and interferon responses. Critically, FRA1 directly activates *AXL*, *CDK6*, and *FSCN1*, which are upregulated in metastatic melanoma and correlate with poor patient outcomes. These findings provide mechanistic insight into how FRA1 promotes metastatic capabilities across multiple steps of the metastatic cascade.

The functional significance of these FRA1 targets was validated through both genetic and pharmacological approaches. Silencing *AXL*, *CDK6*, or *FSCN1* significantly reduced FRA1-mediated invasion and metastatic colonization. Pharmacologically, inhibitors targeting CDK4/6 and Fascin, and to a lesser extent AXL, suppressed metastasis in experimental models. The combination of CDK4/6 and Fascin inhibitors demonstrated particular efficacy in suppressing metastasis and extending survival. These results not only validate our mechanistic findings but also identify potential therapeutic strategies for FRA1-high metastatic melanoma.

Our findings align with studies in other cancer types (6,15), highlighting the role of FRA1 in promoting metastasis. FRA1 promotes metastatic progression of triple negative breast cancer (reviewed in (16)), pancreatic ductal adenocarcinoma (17,18), colon adenocarcinoma (19,20), squamous cell carcinoma (21), and lung adenocarcinoma (22) in experimental models, and a role for FRA1 in influencing EMT as part of the metastatic process has been proposed for several cancers (16,17,20). In melanoma, the FRA1-regulated transcriptional program appears to influence processes related to EMT, including cell migration, adhesion, and ECM interactions. Moreover, MAPK pathway activation induces FRA1 in melanoma cells, which promotes transcription factor reprogramming favoring ZEB1 and TWIST1 expression (23), transcription factors that induce EMT-like states. Accordingly, AXL expression strongly correlates with EMT and focal adhesion signatures (Supplementary Fig. S5B). Additionally, FRA1-mediated repression of apoptosis and the p53 pathway may protect against anoikis, while crosstalk with MYC (24), IL-6 (25), and TGFβ (18) signaling may promote metastasis formation. FRA1 may also suppress interferon responses, and we observed an inverse correlation between *FSCN1* expression and interferon response gene signatures (Supplementary Fig. S5C). Fascin inhibition has been reported to increase intratumoral dendritic cell activation and anti-cancer immunity (26), and these findings suggest possible influences of FRA1 on anti-tumor immunity that warrant further investigation in immune-competent models.

Among the FOS family members, FRA1 appears to play a unique oncogenic role in melanoma. Depletion studies using the DepMap database revealed that silencing FRA1, but not other FOS family members (FOS, FOSB, FRA2), significantly reduces cell viability in melanoma and across multiple cancer types (Supplementary Fig. S5D). This suggests that FRA1 possesses distinct oncogenic functions not shared by its family members, potentially due to unique protein interactions or regulatory mechanisms. FRA1 functions as an obligate heterodimeric transcription factor classically partnering with JUN proteins to form the AP-1 complex. However, our motif analysis revealed enrichment of binding sites for additional transcription factors, including TEAD, ELK1/ELK4, and RUNX, suggesting possible co-regulatory relationships. Previous studies have demonstrated that TEAD and AP-1 can coordinate transcriptional activity in various cancer cell lines (27). Future studies will define the specific transcription factor complexes containing FRA1 that drive melanoma progression, as these interactions may represent additional therapeutic vulnerabilities.

The PTEN/AKT/mTORC1/FRA1 signaling axis we previously identified (5) now extends to the regulation of *AXL*, *CDK6*, and *FSCN1*, suggesting that PTEN loss promotes melanoma metastasis partly through this mechanism. Interestingly, AXL and Fascin have been implicated in reciprocal regulation with AKT, indicating complex regulatory networks controlling melanoma progression. AXL functions as an upstream regulator of AKT-mediated resistance to BRAF inhibitors in melanoma with wildtype PTEN (28), while AKT and Fascin regulate each other (29,30), creating potential feedback mechanisms that may stabilize the pro-metastatic state.

The widespread relevance of the FRA1 transcriptional network is evident across multiple aggressive cancer types, where *FRA1* expression correlates with *AXL*, *CDK6*, and *FSCN1* levels and poor patient outcomes. This conserved signature suggests common mechanisms driving metastasis across distinct cancer types. Our findings establish FRA1 as a potent driver of an actionable pro-metastatic transcriptional network in melanoma and potentially other cancers, offering new avenues for therapeutic intervention in advanced disease with otherwise limited treatment options.

## MATERIALS AND METHODS

### Cell lines and culture conditions

Mouse melanoma cell line M10M6 was established from a tumor from a Braf^V600E^; Pten^null^ (BPP) mouse as described before (31,32). Human melanoma cell line 1205Lu (RRID:CVCL_5239) was provided by M. Herlyn (Wistar Institute, Philadelphia, PA). A375 (RRID: CVCL_6233) and HEK293T Lenti-X were obtained from ATCC and Takara, respectively. All cell lines were cultured in a humidified atmosphere at 37°C and 5% CO2. Melanoma cell lines were cultured in RPMI-1640 (Lonza) containing 5% FBS (Sigma). HEK293T Lenti-X were cultured in DMEM (VWR) containing 10% FBS. All cell lines were confirmed bimonthly to be Mycoplasma-free by the MycoAlert Mycoplasma Detection Kit (Lonza) and STR authenticated when they were initially obtained. Cell lines were used for experiments within 15 passages after thawing.

### Plasmid and lentivirus production

pLenti-CMV-FOSL1-Hygro was generated by replacing GFP in pLenti-CMV-GFP-Hygro with the FOSL1 cDNA from human A375 melanoma cells. The TET-dCas9 vector was generated using Takara In-Fusion cloning (Cat# 639650). The dCas9 sequence was PCR-amplified with 2X Platinum SuperFi II DNA Polymerase from the dCas9-KRAB plasmid (Addgene #110820) and subsequently cloned into the pRRL-Blasticidin vector. pLenti-CRISPRv2-puro was obtained from Addgene (#52961). The Cas9 coding sequence was removed from pLenti-CRISPRv2-puro to create a vector exclusively for short guide RNA (sgRNA) expression. Modifications of the sgRNA sequences were performed using the Q5 Site-Directed Mutagenesis Kit according to the manufacturer’s instructions. sgRNA sequences are listed in Supplementary Table S1. To produce lentivirus supernatants, HEK293T cells were transfected with lentiviral vector and Δ8.2 and VSVG helper plasmids at a 9:8:1 ratio. Lentiviral supernatant was cleared using 0.45 µm syringe filters and used to infect melanoma cells in the presence of 8 mg/mL polybrene. Infected cells were selected with the appropriate antibiotics (1 mg/mL puromycin, 100 mg/mL hygromycin, or 10 mg/mL blasticidin).

### Animal models

All animal experiments were conducted in accordance with an IACUC protocol approved by the University of South Florida. NSG mice were obtained from JAX (Stock No: 005557) and bred in-house. 6-week-old male and female NSG mice were randomly divided into groups (at least 5 mice per group). 200,000 A375 and 1205Lu melanoma cells were subcutaneously injected into the flank of NSG mice. Mice were fed chow containing 200 mg/kg Doxycycline (Envigo). Tumors were measured using calipers and volume was calculated using the formula (width^2^ x length)/2, and spontaneous lung metastases were quantified at endpoint by H&E staining. 200,000 1205Lu and M10M6 melanoma cells were intravenously injected into NSG mice via the tail vein, and lung metastasis burden was monitored after 18-22 days by IVIS in vivo bioluminescence imaging. For IVIS imaging, mice were anesthetized with isoflurane and then intraperitoneally injected with 100 µL Luciferin (4 mg/ml). Bioluminescent imaging was performed 10 min following Luciferin injection using the IVIS Spectrum (Caliper Life Sciences). Mice were euthanized after IVIS imaging, and lungs and livers were resected and fixed in Formalin. Tissue embedding in paraffin, sectioning, and H&E staining were performed by IDEXX. For experimental metastasis models, NSG mice were intravenously inoculated with 200,000 A375 and 1205Lu melanoma cells and euthanized after 7 days (initial colonization) or 24 days (outgrowth). For outgrowth experiments, mice were fed Doxycycline chow (200 mg/kg) from day 10 to day 24 and lung metastasis burden was monitored by IVIS imaging. Lungs and livers were resected and stained with H&E. The number and size of metastatic colonies were quantified by Aperio ImageScope and QuPath.

### Inhibitor treatment

Bemcentinib, Abemaciclib, G1T38, and NP-G2-044 were purchased from Selleckchem and dissolved in DMSO at 12.5 mg/ml, aliquot into 20 μl vials, and stored at -20°C. Before administration, inhibitors were thawed at room temperature and diluted 1:10 with 180 μl sterile PBS. 250,000 1205Lu melanoma cells were intravenously inoculated into NSG mice. After 14 days, mice were treated with Bemcentinib, G1T38, or NP-G2-044 at a dose of 250 µg per mouse, or treated with vehicle control by intraperitoneal (IP) administration every other day for a total of 4 administrations. To test the combination of targeting CDK6 and Fascin, NSG mice were intravenously inoculated with 500,000 1205Lu melanoma cells, and immediately treated with 500 μg Abemaciclib plus 250 µg NP-G2-044, or vehicle control via IP administration daily until endpoint. Metastasis was monitored by CT scan at day 10 and day 18. Endpoint of each mouse was determined based on the metastatic burden from CT scan and the health condition (10% weight loss, labored breathing, hunched posture), and the overall survival was recorded.

### RNA sequencing

RNA extracted from A375 and 1205Lu cells was quantified with a Qubit Fluorometer (ThermoFisher Scientific, Waltham, MA) and screened for quality on the Agilent TapeStation 4200 (Agilent Technologies, Santa Clara, CA). The samples were then processed for RNA sequencing using the NuGEN Universal RNA-Seq Library Preparation Kit with NuQuant (Tecan Genomics, Redwood City, CA). Briefly, 100 ng of RNA were used to generate cDNA and a strand-specific library following the manufacturer’s protocol. Quality control steps were performed, including TapeStation size assessment and quantification using the Kapa Library Quantification Kit (Roche, Wilmington, MA). The final libraries were normalized, denatured, and sequenced on the Illumina NovaSeq 6000 sequencer with the S1–200 reagent kit to generate approximately 60 million 100-base read pairs per sample (Illumina, Inc., San Diego, CA). Read adapters were detected using BBMerge (v37.02) (33) and subsequently removed with cutadapt (v1.8.1) (34). Processed raw reads were then aligned to human genome GRCh38 using STAR (v2.7.7a) (35). Gene expression was evaluated as read count at gene level with RSEM (v1.3.0) (36) and Gencode gene model v30. Gene express data were then normalized and differential expression between experimental groups were evaluated using DEseq2 (37).

### Cut&Run sequencing

Cut&Run (38) was performed with A375 and 1205Lu cells using CUTANA ChIC/CUT&RUN Kit version 4.0 (EpiCypher, Cat. # SKU: 14–1048). Briefly, 500,000 cells were collected and re-suspended in Wash Buffer and mixed with 10 μl activated ConA beads. 0.5 μg FRA1 antibody (Abcam, Cat. # ab252421) or 1 μl IgG antibody were added and incubated overnight at 4°C. Beads were washed twice in 200 μl Cell Permeabilization Buffer (CPB) prior to the addition of 2.5 μl CUTANA pAG-MNase in 50 μl of CPB per reaction. Beads were again washed twice in 200 μl CPB and suspended in 50 μl CPB. For chromatin digestion, 1 μl of 100 mM CaCl_2_ was added to activate the MNase. 33 μl of Stop Master Mix was added to terminate the MNase activity after 2h incubation at 4°C. To isolate DNA from the supernatant, 119 μl SPRlselect were added and incubated for 5 minutes at room temperature. SPRIselect beads were washed twice with 85% Ethanol and air-dried. To elute DNA, 17 μl of 0.1X TE Buffer were added. Eluted DNA was quantified with the Qubit fluorometer, and 0.6 to 5 nanograms of enriched DNA was used for library preparation using the Kapa HyperPrep Kit (Roche, Cat. # 07962347001) following EpiCypher’s parameters for indexing PCR and library amplification as described in the CUT&RUN Library Prep Manual. The final libraries were Qubit quantitated and screened on the Agilent TapeStation D1000 ScreenTape (Agilent Technologies) to assess the fragment size distribution. Following final quantification using the Kapa Library Quantification Kit (Roche, Cat. # 07960140001), the libraries were sequenced on the NextSeq 500 using a Mid-output 150-cycle Kit in 2×50 configuration to generate 6–8 million read-pairs per sample (Illumina). Read adapters were detected using BBMerge (v37.02) (33) and subsequently removed with cutadapt (v1.8.1) (34). Processed raw reads were then aligned to human genome GRCh38 using BOWTIE2 (v2.4.1) (39). Peak Calling was performed to identify regions of open chromatin using MACS2 (v 2.1.1.20160309). Only peaks significantly enriched over background with q <0.01 were considered confident and subjected to downstream analysis. Peaks were annotated by ChIPseeker (v1.32.1) (40) with gene model TxDb.Hsapiens.UCSC.hg38.knownGene (v 3.15.0). Consensus peak identifiers were obtained by GenomicRanges (v1.48.0) (41). Peaks were then normalized and differential enrichment between experimental groups evaluated using DEseq2 (v1.36.0) (42).

### TCGA data and CCLE data

Gene expression data and patient outcome data from the following cancer types in The Cancer Genome Atlas (TCGA) Pan Cancer Atlas were extracted from cBioPortal (43) and analyzed for the various correlation and survival studies using Kaplan–Meier analysis: Skin cutaneous melanoma, Uveal melanoma, Lung adenocarcinoma, Pancreatic adenocarcinoma, Kidney renal clear cell carcinoma, Sarcoma, Low grade glioma, Glioblastoma, Head and Neck squamous cell carcinoma, Colon adenocarcinoma, Bladder urothelial carcinoma, Liver hepatocellular carcinoma, Prostate adenocarcinoma, Lung squamous cell carcinoma, Stomach adenocarcinoma, Breast invasive carcinoma, Esophageal carcinoma (ESCC). Gene expression data from the following cancer types in Cancer Cell Line Encyclopedia (CCLE) dataset (44,45) were extracted from DepMap Portal (46) and analyzed for the correlation between *FOSL1* expression and expression of *AXL*, *CDK6*, and *FSCN1*: Breast invasive carcinoma, Bladder urothelial carcinoma, Non-small cell lung cancer, Colon adenocarcinoma, Head and Neck squamous cell carcinoma, Esophageal carcinoma, Stomach adenocarcinoma, Prostate adenocarcinoma, Kidney renal clear cell carcinoma.

### Gene set enrichment analysis

RNA sequencing data of A375 and 1205Lu melanoma cells were subjected to gene set enrichment analysis (GSEA; RRID:SCR_003199) using “Hallmark gene set”, “KEGG_legacy subset of Canonical pathways”, and a melanoma metastasis gene signature “Winnepenninckx_Melanoma_Metastasis_Up” to characterize the FRA1 transcriptome. “si-NT” and “si-FRA1” were used as phenotype, and “No_Collapse” was used for gene symbol. The metric for ranking genes in GSEA was set as “Signal2Noise”, otherwise default parameters were used. Global mRNA expression profiles of TCGA skin cutaneous melanoma dataset (TCGA-SKCM) were subjected to GSEA to identify the association of *FOSL1* with gene signature “Winnepenninckx_Melanoma_Metastasis_Up”. Expression of *FOSL1* was used as phenotype, and “No_Collapse” was used for gene symbol. The metric for ranking genes in GSEA was set as “Pearson,” otherwise default parameters were used. GSEA was performed using GSEA 4.3.3 software.

### Proliferation assay

Four thousand 1205Lu cells/well were seeded in 96 well plates in 200 µL RPMI-1640 medium containing 5% FBS. After 24 hours, plates were loaded into Cellcyte-X live cell analyzer (ECHO). Images of each well were taken daily for 6 continuous days and the cell confluency of each image was quantified by the Cellcyte analyzer software.

### Transwell invasion assay

To pre-coat transwells, Matrigel (Corning) was thawed on ice for at least 2 hours, diluted to 5% with ice-cold RPMI-1640 medium, gently added to transwell inserts (50 µL/insert), and solidified at 37°C for 30 min. 40,000 1205Lu cells were plated per Matrigel-coated insert in 200 µl of RPMI-1640 without FBS. 500 µl RPMI-1640 medium containing 15% FBS were added into the bottom well. Plates were incubated at 37°C in a humidified incubator for 48 hr. Media was discarded, inserts were gently washed once with PBS, and cells were fixed with 500 µl fixing solution (ethanol:acetic acid = 3:1) for 10 minutes. Inserts were washed once with PBS, and cells were stained in 500 µL staining solution (0.5% crystal violet) for 30 minutes. Inserts were washed with tap water twice and non-migrated cells and Matrigel on the top side of the insert were carefully removed with cotton swabs, and inserts were air dried overnight. Pictures were taken at 100x magnification and cell numbers quantified.

### qRT-PCR

RNA was extracted from cells using TRIzol (Invitrogen) following protocols supplied by the manufacturer. cDNA was generated with PrimeScript RT Master Mix (Takara). qPCR was performed on StepOnePlus Real-Time PCR System (Thermo Fisher Scientific) using PerfeCTa SYBR Green Fastmix (Quantabio). mRNA expression was normalized to 18S rRNA. Primer sequences were obtained from PrimerBank (https://pga.mgh.harvard.edu/primerbank/index.html; RRID:SCR_006898) and are listed in Supplementary Table S1.

### Western blot analysis

15 µg protein were separated on NuPAGE 4-12% precast gels (Thermo Fisher Scientific) and transferred to nitrocellulose membranes. Membranes were blocked in 5% non-fat dry milk in TBST and incubated with one of the following primary antibodies overnight at 4°C: FRA1 (1:1,000), AXL (1:1,000), CDK6 (1:1,000), Fascin (1:1,000). Anti-beta-actin (1:3,000) was used as loading control. Membranes were washed 3 times with TBST for 10 min, followed by incubation with HRP-conjugated secondary antibodies (1:3,000) for 1 hr at room temperature. After three washes in TBST, chemiluminescence substrate (1:1) was applied to the blot for 4 min and chemiluminescence signal was captured using a LI-COR imaging system.

### Tissue Microarray staining

A melanoma TMA was stained for FRA1, AXL, CDK6, Fascin, and DAPI with an Automated Opal 7-Color IHC Kit and quantified in Moffitt’s Advanced Analytical and Digital Laboratory:

*Multiplex Immune Panel Procedure:* Formalin-fixed and paraffin-embedded (FFPE) tissue samples were immunostained using the AKOAYA Biosciences OPAL 7-Color Automation IHC kit (Waltham, MA) on the BOND RX autostainer (Leica Biosystems, Vista, CA). The OPAL 7-color kit uses tyramide signal amplification (TSA)-conjugated to individual fluorophores to detect various targets within the multiplex assay. Sections were baked at 65°C for one hour and then transferred to the BOND RX (Leica Biosystems). All subsequent steps (ex., deparaffinization, antigen retrieval) were performed using an automated OPAL IHC procedure (AKOYA). OPAL staining of each antigen occurred as follows: heat induced epitope retrieval was achieved with EDTA pH 9.0 buffer for 20 min at 95°C before the slides were blocked with AKOYA blocking buffer for 10 min. Then slides were incubated with primary antibody at RT for 30 min followed by OPAL HRP polymer and one of the OPAL fluorophores during the final TSA step. Individual antibody complexes are stripped after each round of antigen detection. This was repeated five more times using the remaining antibodies of the panel. After the final stripping step, DAPI counterstain was applied to the multiplexed slide and was removed from BOND RX for coverslipping with ProLong Diamond Antifade Mountant (ThermoFisher Scientific). Autofluorescence slides (negative control) were included, which used primary and secondary antibodies omitting the OPAL fluorophores and DAPI. All slides were imaged with the Phenolmager HT (Akoya Biosciences). *Quantitative Image Analysis*: Multi-layer TIFF images were exported from InForm (AKOYA) and loaded into HALO Image Analysis Platform version 4.0 (Indica Labs, New Mexico) for quantitative image analysis. For the quantitative fluorescent phenotype analysis, tissues were segmented into individual cells using the DAPI marker. For each marker, a positivity threshold within the nucleus or cytoplasm is determined based on visual intensity and expected staining localization. After setting a positive fluorescent threshold for each staining marker, the entire image set is analyzed with the created algorithm. The generated data includes positive cell counts for each fluorescent marker in cytoplasm or nucleus, and percent of cells positive for the marker.

### Quantification and statistical analysis

Proliferation, low density growth assay, and soft agar assay are presented as mean ± SEM. For transwell invasion assays, 3-4 random fields were quantified for each well and data are presented as mean ± SEM. All experiments were performed at least three times with three to four technical replicates. Statistical analyses were performed by Student’s unpaired t test. Subcutaneous tumor growth curves and tumor weights at endpoint are presented as mean ± SEM. Graphpad Prism 10 (RRID:SCR_002798) was used for statistical analyses and a P value of <0.05 was considered statistically significant.

### Data availability

Sequencing data supporting the findings of this study have been deposited in the Gene Expression Omnibus under the accession codes GSE293767 (https://www.ncbi.nlm.nih.gov/geo/query/acc.cgi?acc=GSE293767; token for access: sncfwugsphortoz). Source data are provided with this paper. All other data supporting the findings of this study are available from the corresponding author upon request.

## ACKNOWLEDGMENTS

We are grateful to Karreth lab members for helpful discussions. X.X. was supported by a Melanoma Research Foundation Career Development Award (1068914). F.A.K. received funding from the NIH/NCI (R01CA259046) and the Florida Department of Health Bankhead Coley Program (21B04). This work was supported by the Molecular Genomics Core, Biostatistics and Bioinformatics Shared Resource, Small Imaging Lab, and the Advanced Analytical and Digital Laboratory which are funded in part by Moffitt’s Cancer Center Support Grant (P30CA076292).

## AUTHORS CONTRIBUTIONS

**X.X.**: Conceptualization, formal analysis, investigation, visualization, methodology, writing–original draft, writing–review and editing. **M.C.**: Investigation. **Z.S-V.**: Investigation. **V.J.**: Investigation. **J.Y.**: Formal analysis. **X.Y.**: Formal analysis, investigation, methodology. **F.A.K.**: Conceptualization, formal analysis, supervision, funding acquisition, investigation, visualization, methodology, writing–original draft, project administration, writing–review and editing.

## SUPPLEMENTARY FIGURE LEGENDS

**Supplementary Figure S1.**
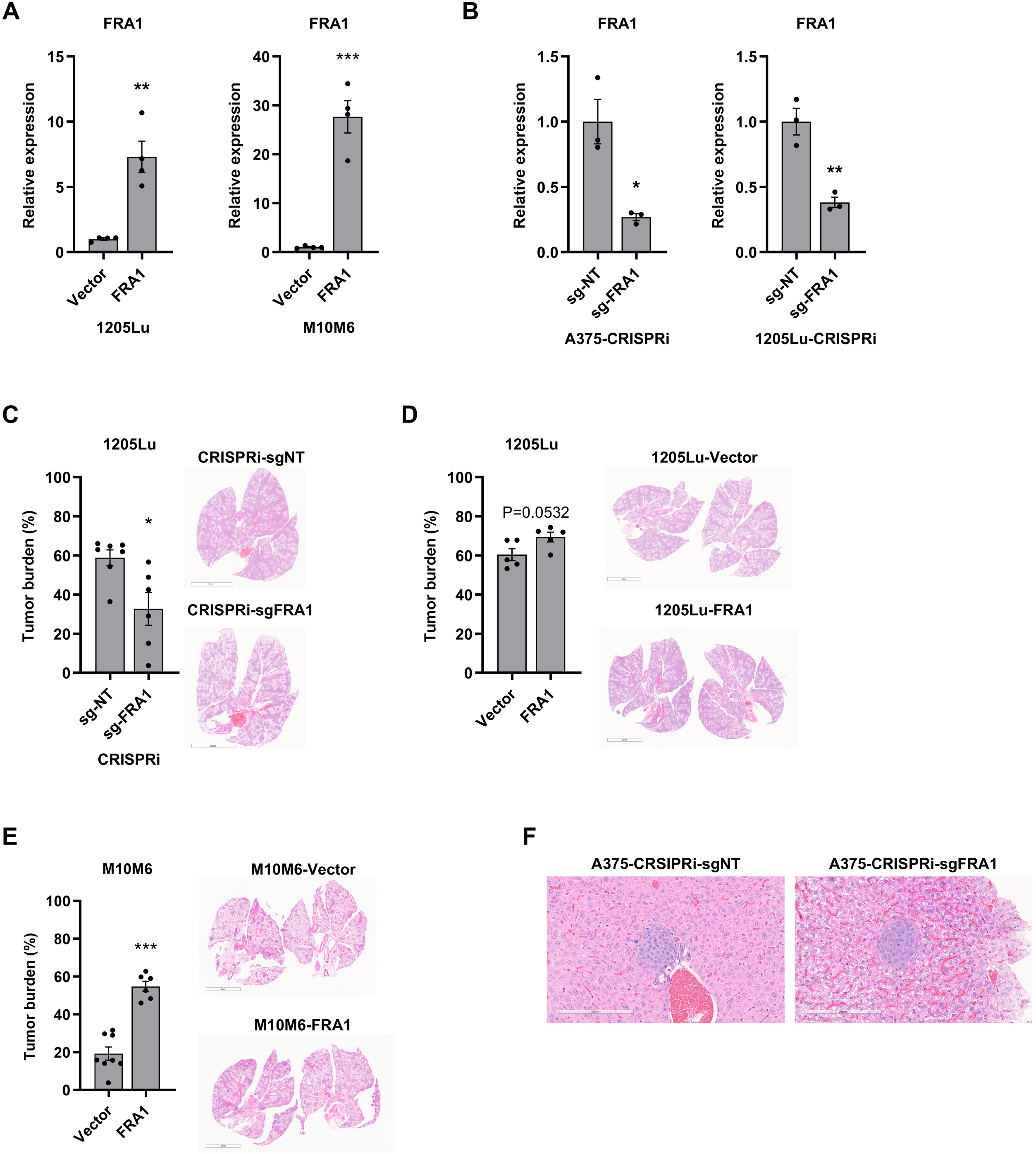
**A,** qRT-PCR show the relative expression of FRA1 in FRA1 overexpressing 1205Lu and M10M6 cells. **B,** qRT-PCR show the relative expression of FRA1 in A375 and 1205Lu cells with CRISPRi targeting FRA1. **C-E,** Luciferase tagged 1205Lu cells with inducible CRISPRi targeting FRA1 were intravenously injected into NSG mice (n = 5) and the metastases were measured after 22 days by H&E staining (**C**). Luciferase tagged 1205Lu and M10M6 cells with FRA1 overexpression were intravenously injected into NSG mice (n = 5), and mice were fed chow containing 200 mg/kg Dox to induce CRISPRi. The metastases were measured after 18 days by H&E staining (**D** and **E**). Representative images of H&E staining and quantification of metastatic burdens are shown. **F,** Luciferase tagged and A375 cells with inducible CRISPRi targeting FRA1 were intravenously injected into NSG mice (n = 5), and 10 days after inoculation mice were switched to chow containing 200 mg/kg Dox to induce CRISPRi, and then metastasis were analyzed 2 weeks after Dox diet feeding, which is 24 days after cell inoculation. Representative images of liver metastasis by H&E staining are shown. Data are presented as mean ± SEM and analyzed with Student’s unpaired t test, * P<0.05, ** P<0.01, *** P<0.001.

**Supplementary Figure S2.**
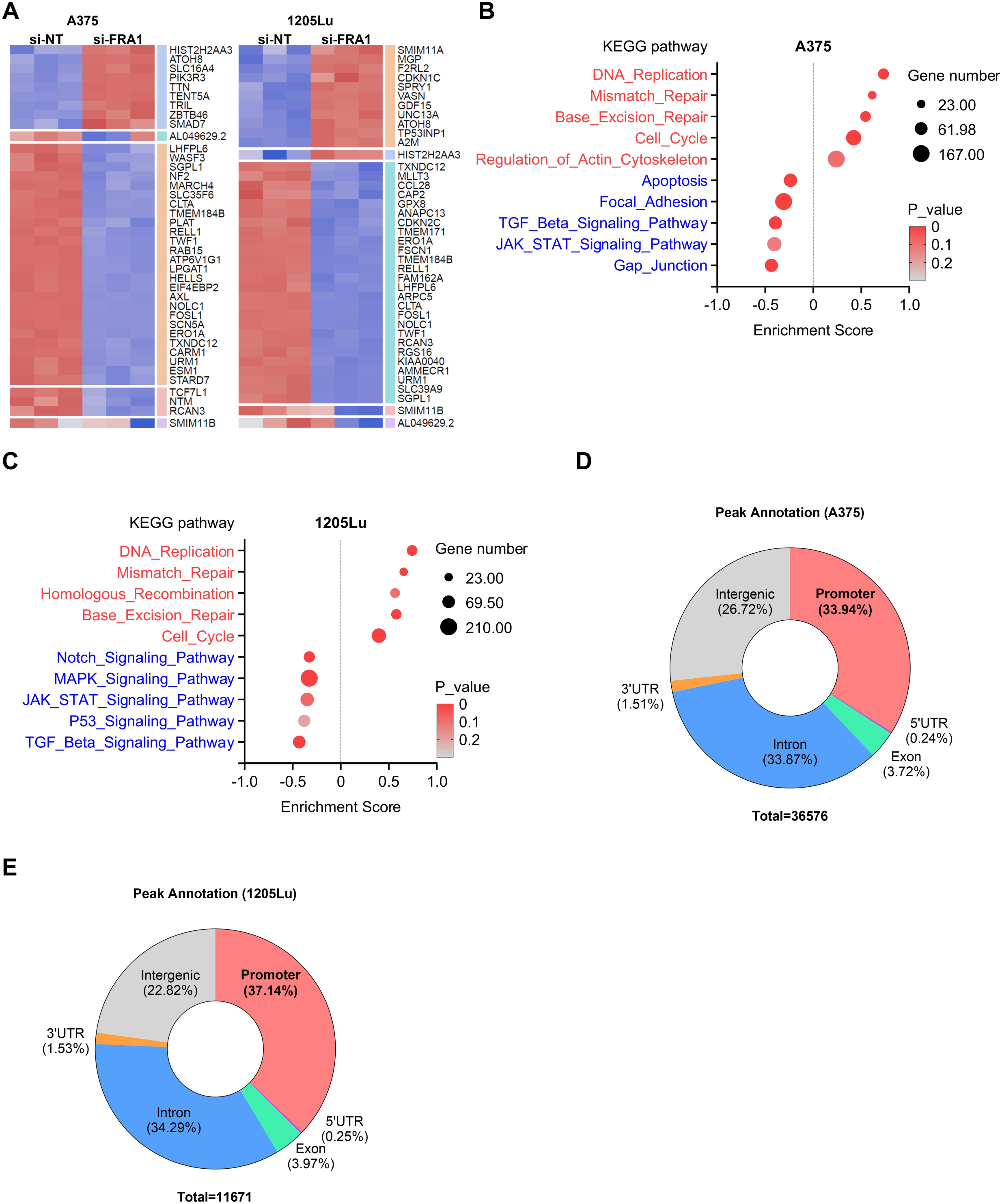
**A,** Heatmap of RNA sequencing showed the top differential expressed genes by silencing FRA1 in A375 and 1205Lu cells. **B, C,** GSEA analysis of KEGG pathways show enriched gene signatures associated with FRA1 transcriptome in A375 cells (**B**) and 1205Lu cells (**C**). **D, E,** Genomic annotation of FRA1 bound peaks in A375 cells (**D**) and 1205Lu cells (**E**).

**Supplementary Figure S3.**
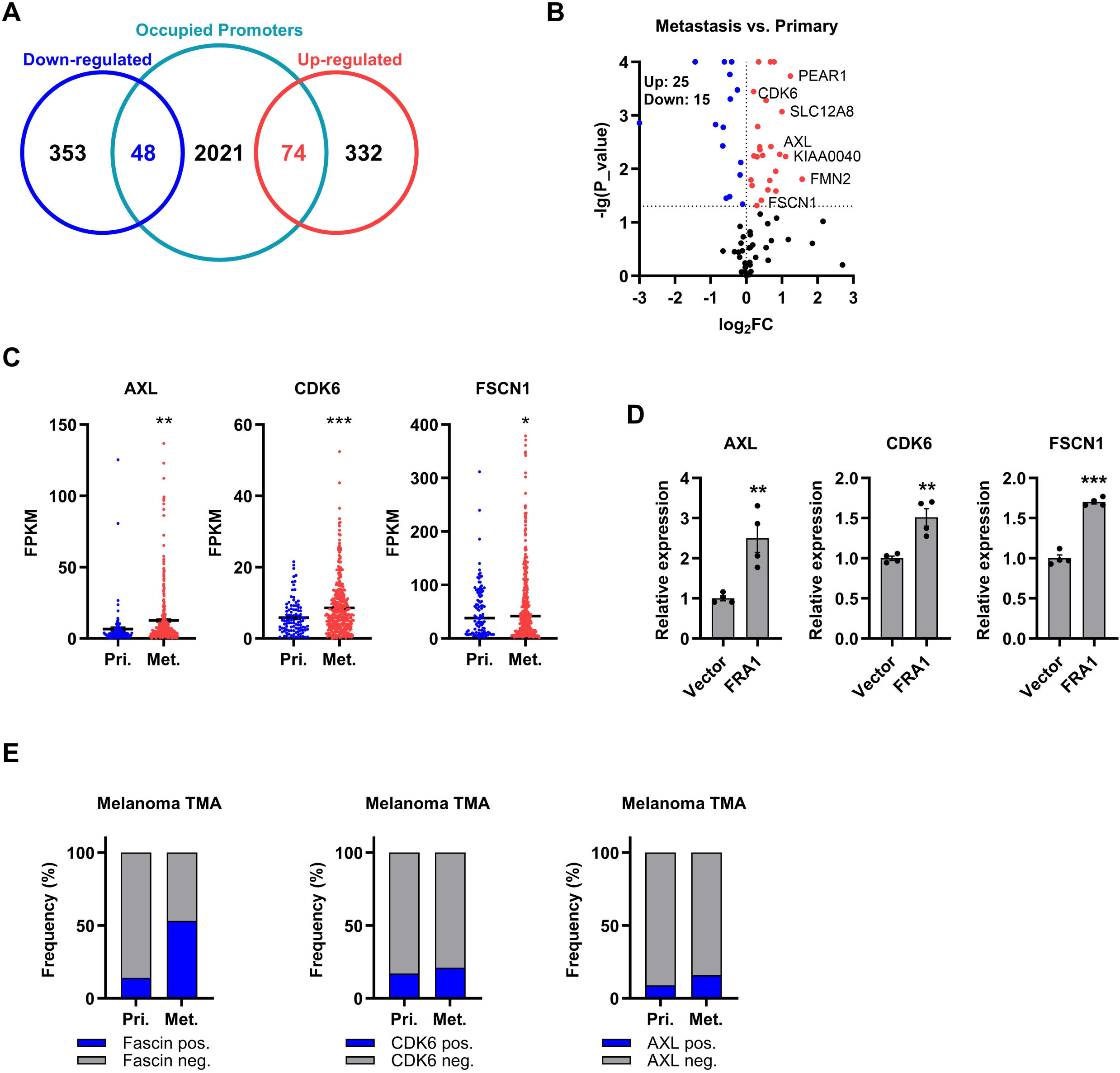
**A,** Venn diagram showing the overlap of differentially expressed genes from RNA sequencing and genes containing FRA1 at promoter regions from Cut&Run sequencing. **B,** A total of 74 genes were identified as potential targets transcriptionally activated by FRA1, and their differential mRNA expression between metastatic melanoma and primary melanoma was analyzed in the TCGA-SKCM dataset and is shown as volcano plot. **C,** Expression of *AXL*, *CDK6*, and *FSCN1* in primary and metastatic melanoma samples in the TCGA-SKCM dataset. **D,** qRT-PCRs showing the relative expression of *AXL*, *CDK6*, and *FSCN1* upon expression of FRA1 in 1205Lu cells. **E,** Bar plots showing the frequency of *AXL*, *CDK6*, or *FSCN1* positive samples in primary skin melanomas and lymph node metastatic melanomas. Data are presented as mean ± SEM and analyzed with Student’s unpaired t test, * P<0.05, ** P<0.01, *** P<0.001.

**Supplementary Figure S4.**
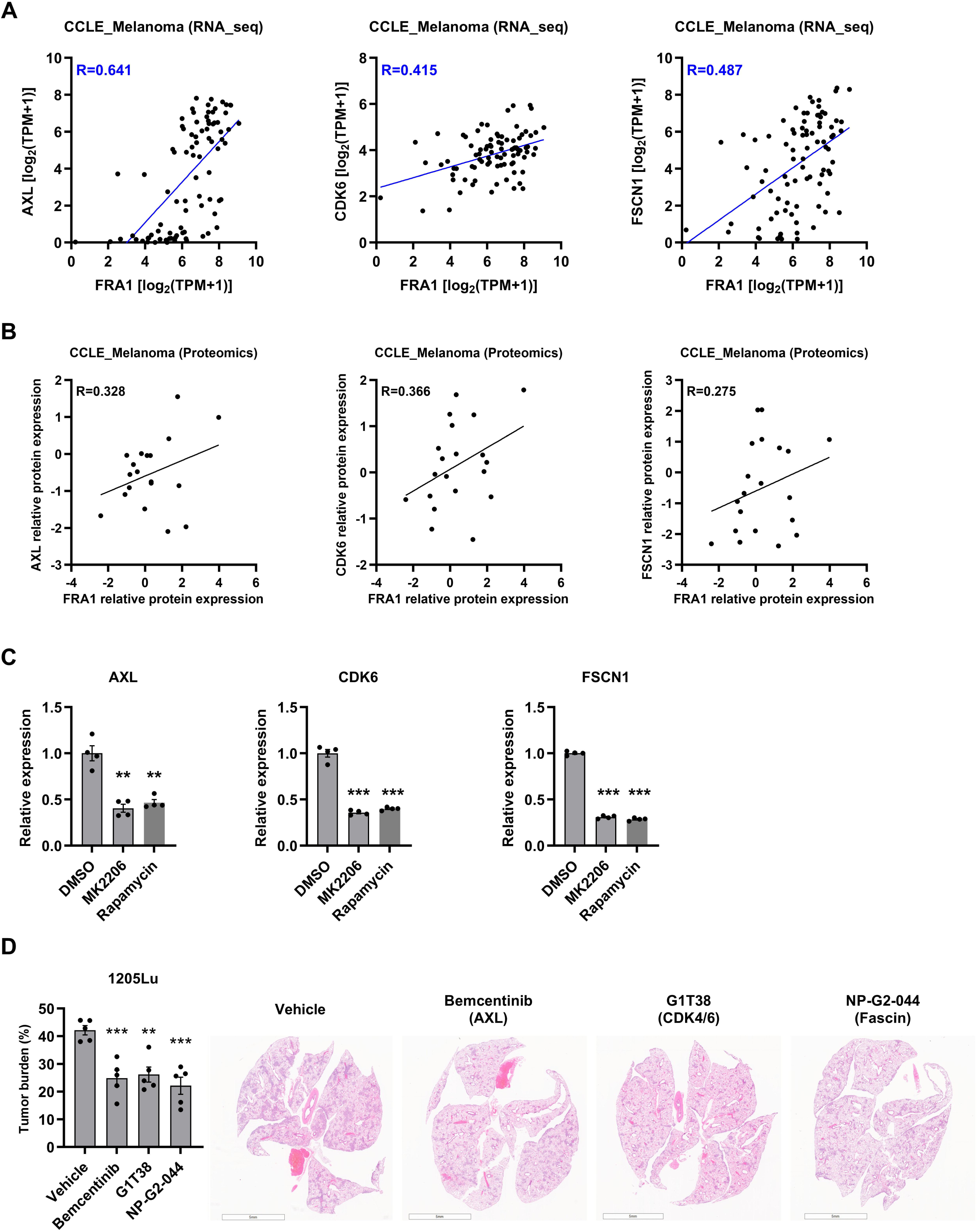
**A,** Correlation of *FRA1* mRNA level and mRNA of *AXL*, *CDK6*, and *FSCN1* in 87 melanoma cell lines from Cancer Cell Line Encyclopedia (CCLE) RNA-seq dataset. **B,** Correlation of FRA1 protein level and protein of AXL, CDK6, Fascin in 20 melanoma cell lines from CCLE Proteomics dataset (data extracted from DepMap). **C**, qRT-PCR showing the relative expression of *AXL*, *CDK6*, and *FSCN1* in 1205Lu cells upon treatment with AKT inhibitor MK2206 or mTOR inhibitor Rapamycin. **D,** Luciferase tagged 1205Lu cells were intravenously injected into NSG mice. AXL inhibitor Bemcentinib, CDK4/6 inhibitor G1T38, Fascin inhibitor NP-G2-044, and vehicle control were intraperitoneally (I.P.) administrated to mice (n = 5) at day 14, 16, 18, and 20 after cell inoculation. The metastases were measured at day 20 by H&E staining. Representative images of H&E staining and quantification of metastatic burdens are shown. Data are presented as mean ± SEM and analyzed with Student’s unpaired t test, * P<0.05, ** P<0.01, *** P<0.001.

**Supplementary Figure S5.**
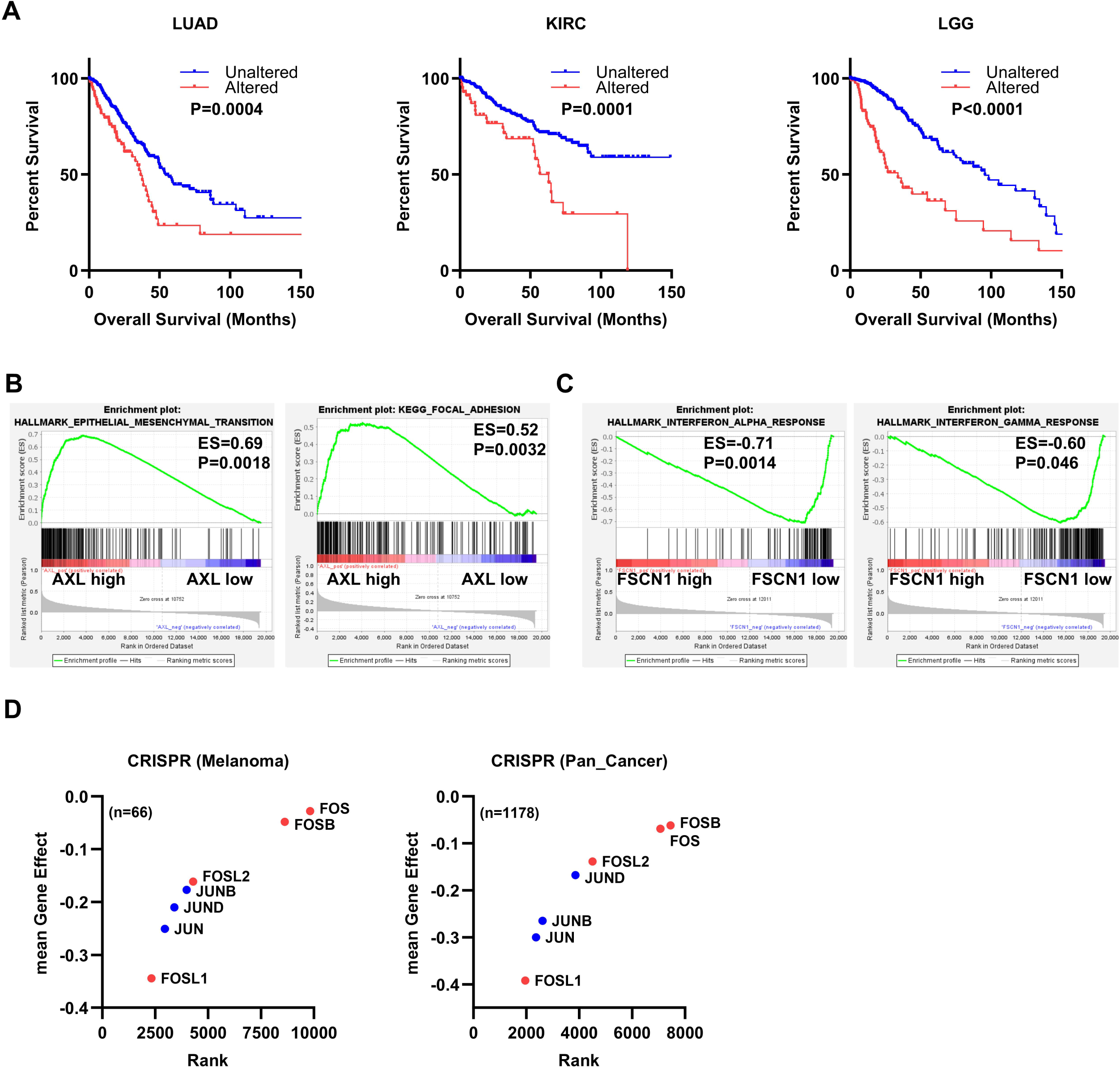
**A,** Overall survival curves of patients with or without alteration of AXL, CDK6, or FSCN1 (including mutation, copy number alteration, and mRNA changes) of multiple cancer types TCGA dataset (data extracted from cBioPortal). **B,** GSEA analysis of TCGA-SKCM samples associates AXL expression with gene signatures of epithelial-mesenchymal transition (Hallmark gene signature) and focal adhesion (KEGG pathway). **C,** GSEA analysis of TCGA-SKCM samples associates FSCN1 expression with gene signatures of interferon alpha and gamma response (Hallmark gene signature). **D,** Gene dependency based on CRISPR knockout screening in 66 melanoma cell lines and 1178 cell lines of all cancer types (data extracted from DepMap).

## Notes

The authors have no conflicts of interest to declare.

### Competing Interest Statement

The authors have declared no competing interest.

